# The association between cortical gyrification and sleep in adolescents and young adults

**DOI:** 10.1101/2023.09.15.557966

**Authors:** João Paulo Lima Santos, Rebecca Hayes, Peter L. Franzen, Tina R. Goldstein, Brant P. Hasler, Daniel J. Buysse, Greg J. Siegle, Ronald E. Dahl, Erika E. Forbes, Cecile D. Ladouceur, Dana L. McMakin, Neal D. Ryan, Jennifer S. Silk, Maria Jalbrzikowski, Adriane M Soehner

## Abstract

**Study objectives:** Healthy sleep is important for adolescent neurodevelopment, and relationships between brain structure and sleep can vary in strength over this maturational window. Although cortical gyrification is increasingly considered a useful index for understanding cognitive and emotional outcomes in adolescence, and sleep is also a strong predictor of such outcomes, we know relatively little about associations between cortical gyrification and sleep.

**Methods:** Using Local gyrification index (lGI) of 34 bilateral brain regions and regularized regression for feature selection, we examined gyrification-sleep relationships in the Neuroimaging and Pediatric Sleep databank (252 participants; 9-26 years; 58.3% female) and identified developmentally invariant (stable across age) or developmentally specific (observed only during discrete age intervals) brain-sleep associations. Naturalistic sleep characteristics (duration, timing, continuity, and regularity) were estimated from wrist actigraphy.

**Results:** For most brain regions, greater lGI was associated with longer sleep duration, earlier sleep timing, lower variability in sleep regularity, and shorter time awake after sleep onset. lGI in frontoparietal network regions showed associations with sleep patterns that were stable across age. However, in default mode network regions, lGI was only associated with sleep patterns from late childhood through early-to-mid adolescence, a period of vulnerability for mental health disorders.

**Conclusions:** We detected both developmentally invariant and developmentally specific ties between local gyrification and naturalistic sleep patterns. Default mode network regions may be particularly susceptible to interventions promoting more optimal sleep during childhood and adolescence.

## INTRODUCTION

Brain structure and sleep patterns undergo significant maturational changes over adolescence [1], and developmental shifts in these phenomena are each known to influence adolescent emotional, social, cognitive, and behavioral outcomes [2–4]. Data from animal models now demonstrates that sleep quality during the sensitive period of adolescence plays a causal role in adult behavioral outcomes via brain-based pathways [5]. Yet, our basic understanding of relationships between brain morphology and sleep patterns in human adolescents remains in the early stages. Although several studies have now explored relationships between gray matter structure (e.g., cortical thickness, cortical and subcortical volume) and sleep in adolescence, we still know relatively little about associations between cortical gyrification and sleep [6]. Cortical gyrification (i.e., the folding of the cerebral cortex [7–9]) is a sensitive indicator of brain development [10, 11] and is emerging as an important predictor of positive adolescent health outcomes [12, 13]. Exploring the relationship between cortical gyrification and sleep during adolescence will deepen our understanding of complex relationships between the brain and sleep during the transition from childhood to adulthood.

In the human brain, gyrification starts in utero, leading to an increase in cortical surface area that fosters a marked increase in the number of neurons and neuronal connectivity without increasing overall brain volume [7–9, 14]. Gyrification peaks approximately two years after birth, and decreases across the lifespan [8, 9] in a non-linear trajectory [9], with childhood and adolescence showing more accelerated decreases over time relative to adulthood [9], most likely due to the maturational changes happening in the adolescent brain [14]. Overall, lower gyrification has been associated with worse outcomes in adults (e.g., poorer memory, attention, executive function) [10, 15]. However, mixed anomalous patterns of cortical gyrification have been reported by studies focusing on major psychiatric disorders [16–20]. Cortical gyrification is related to other structural MRI measures (e.g., volume, thickness, and surface area) [7, 15]; however, these measures only partially overlap [7]. Thus, cortical gyrification may provide unique information about brain-sleep relationships across neurodevelopment.

Given the importance of sleep for many aspects of healthy development (e.g., mental health, cognition, emotional regulation) [4, 21, 22], understanding whether the relationship between cortical gyrification and sleep is developmentally invariant (stable across age) or developmentally specific observed only during discrete age intervals or windows of development) may offer new insights into windows of vulnerability to brain-based risk and windows of opportunity for early intervention promoting healthy development. We recently showed that, between 9 and 26 years, regional cortical thickness and subcortical volumes have both developmentally invariant and developmentally specific (relationships with naturalistic sleep patterns (sleep duration, timing, continuity, and regularity) [1]. These findings suggest that neural mechanisms that underlie brain-sleep relationships have both stable and dynamic features [1], findings which have implications for targeted interventions and for understanding neurobiological mechanisms of sleep. However, given the unique developmental trajectory of cortical gyrification, particularly in comparison to thickness and subcortical volumes [7], it is unclear whether gyrification will also exhibit developmentally specific associations with sleep during adolescence. Our goal was to identify the relationships between cortical gyrification and actigraphy-assessed sleep characteristics in participants between 9-26 years, and to identify whether these relationships are stable over time (developmentally invariant) or present only during specific windows of development (developmentally specific). We used group-lasso regression [23, 24], an exploratory feature selection approach, to identify associations between regional gyrification and four key components of multidimensional sleep health (sleep duration, timing, continuity, and regularity) [2]. We hypothesized that cortical gyrification would show both developmentally invariant and developmentally specific associations with sleep, and that the latter would be more common during periods of increased brain development (childhood and/or adolescence.

## METHODS

### Participants

The NAPS databank includes nine University of Pittsburgh studies that were conducted between 2009 and 2020 and includes baseline actigraphy sleep and structural MRI scan data for all participants (Supplemental Table 1). This databank was approved as a secondary data analysis protocol by the University of Pittsburgh Institutional Review Board. The initial sample in the databank consisted of 394 participants (for additional information about the inclusion criteria, see Supplemental Material). For these analyses, participants were selected if they aged between 9-26 years old, had sleep and neuroimaging data available, good quality MRI, and no history of psychiatric disorders or psychotropic medications. The inclusion of participants between 9-26 years takes into account the broad definition for adolescence (10-24 years) [25]. The final sample included 252 (mean age [SD]= 17.37[4.53], age range=9-26 years, 58.3% female) participants. Demographic and sleep information are detailed in Table 1.

**Table 1.** Demographic and sleep characteristics of the sample included in the analyses.

| **Variable** | **Mean or N (SD or %)** |
| --- | --- |
| Sample *N* | 252 |
| Age (years) | 17.37 (4.53) |
| Self-reported sex |  |
| Female | 147 (58.3%) |
| Male | 105 (41.7%) |
| Ethnicity |  |
| Non-Hispanic | 236 (93.65%) |
| Hispanic | 14 (5.56%) |
| Missing | 2 (0.79%) |
| Race |  |
| White | 181 (71.83%) |
| Black | 38 (15.08%) |
| Asian | 9 (3.57%) |
| Multiple | 21 (8.33%) |
| Unknown/missing | 3 (1.19%) |
| Actigraphy model |  |
| AMI Octagonal MotionLogger | 35 (13.89%) |
| PR/MiniMitter Actiwatch64 | 25 (9.92%) |
| PR Actiwatch2 | 98 (38.89%) |
| PR Spectrum Series | 94 (37.30%) |
| Tracking days | 6.61 (0.83) |
| Weekdays | 4.53 (0.91) |
| Weekend days | 2.07 (0.51) |
| Season |  |
| Spring | 44 (17.46%) |
| Summer | 106 (42.06%) |
| Fall | 58 (23.02%) |
| Winter | 44 (17.46%) |
| Sleep duration (minutes) | 420.54 (62.15) |
| Wake after sleep onset (minutes) | 56.45 (26.87) |
| Midsleep (minutes from midnight) | 265.68 (76.44) |
| Midsleep variability (minutes) | 64.81 (26.87) |

Abbreviations: AMI = Ambulatory Monitoring Incorporated, PR=Philips Respironics

### Sleep data

#### Wrist actigraphy

To behaviorally assess sleep, we used wrist actigraphy, a well-validated method for assessing naturalistic sleep characteristics in youth and adults [26, 27]. Participants wore actigraphs on their non-dominant wrist during a period of five or more consecutive days [28] and were asked to indicate the start and end of each sleep interval via button press. Wrist activity was sampled in 1-minute intervals (epochs). A combination of validated brand-specific sleep algorithms (Philips Respironics-Medium Threshold; Ambulatory Monitoring Incorporated-Sadeh) and standardized visual editing procedures [29, 30] was used to estimate sleep characteristics. For additional information about brands, scoring procedures, and quality assurance, see Supplemental Material, Supplemental Table 2 or [1].

#### Sleep characteristics

Four sleep characteristics were calculated based on actigraphic sleep data: (1) sleep duration (total sleep time in minutes), (2) timing (midpoint between sleep onset and offset in minutes from midnight), (3) continuity (minutes awake after sleep onset; WASO), and (4) regularity (intra-individual standard deviation of midpoint in minutes). The first three characteristics were averaged over the 5–7 tracking days most proximal to their MRI scan; regularity was calculated from the available days of recording. Sleep characteristics (duration, timing, continuity, and regularity) were natural log transformed to normalize distributions.

### Neuroimaging data

Acquisition protocols for each study are described in Supplemental Table 3. Data usability was evaluated using an automated MRIQC T1w-classifier that determined individual scan quality based on a reference template [31]. T1 data were preprocessed using the FreeSurfer analysis software (v6.0) [32–35]. We implemented an additional quality assessment pipeline developed and used by the Enhancing Neuroimaging Genetics through Meta-Analysis consortium [36–46].

Local gyrification index (lGI) was calculated using Freesurfer (-localGI flag; https://surfer.nmr.mgh.harvard.edu/fswiki/LGI). In this approach, local measurements of cortical gyrification were calculated using the ratio between the pial cortical surface (interface between the brain and meninges) and the perimeter of an outer smoothed surface tightly wrapping the pial surface [47]. Brain regions with increased folding show higher lGI, while brain regions with less folding show lower lGI. Given that we did not have any specific hypotheses about laterality, we averaged cortical gyrification across hemispheres for each region (Desikan–Killiany atlas) [48], generating 34 individual measures. To account for scanner effects, we used ComBat, a batch-effect correction tool that minimizes the variance introduced by different MRI protocols [49].

### Statistical approach

To explore the association between cortical gyrification and sleep, we used the R package Group-LassoINTERaction-NET (GLINTERNET) [23, 24] to examine the main relationships between structural neuroimaging measures (lGI), as well as their interaction with age and sex, for each sleep characteristic (sleep duration, timing, continuity, and regularity). Group-lasso is a feature-selection method that identifies the strongest variables associated with an outcome and uses a shrinkage parameter to reduce the coefficient of non-relevant variables toward zero [23, 24]. In addition, this method allows for the inclusion of interactions between non-zero coefficients as potential predictors [24]. We also included multiple actigraphy covariates (i.e., tracking days, season, ratio of weekday to weekend days, actigraph model) in the models. Similar to previous work [1], we repeated 10-fold cross validation 100 times, using the penalty parameter (λ) one standard deviation away from the minimal cross-validation error (Lmin). The final model for each sleep characteristic was the Lmin model that was selected most often during cross validation.

To gain a better understanding of the final model, we followed up on main relationships identified in the above group-lasso models for each sleep characteristic (sleep duration, timing, continuity, and regularity) with multiple regression models. We computed R^2^ estimate variance explained by each full model, as well as groups of measures (i.e. demographics, neuroimaging measures) [50–52]. We then assessed non-zero interactions between age and neuroimaging measures using the Johnson-Neyman technique [53–55], which obtains parameter estimates and points of significance from the interaction between two continuous variables. In this study, the Johnson-Neyman technique helped identify the discrete age intervals (or windows) in which cortical gyrification and sleep components relationships are significant. Non-zero interactions between sex and neuroimaging measures were probed by comparing estimated marginal means [56]. Figures were generated using the R packages ggseg and ggplot2 [57, 58].

## RESULTS

All non-zero associations (main effects and their interactions) for the four sleep characteristics are described in Table 2A-D. The relationship between cortical gyrification and sleep characteristics is shown in Figure 1. The stability of non-zero predictors and multiple regression models are detailed in Supplemental Tables 4 and 5. Each model is described below.

**Figure 1.**
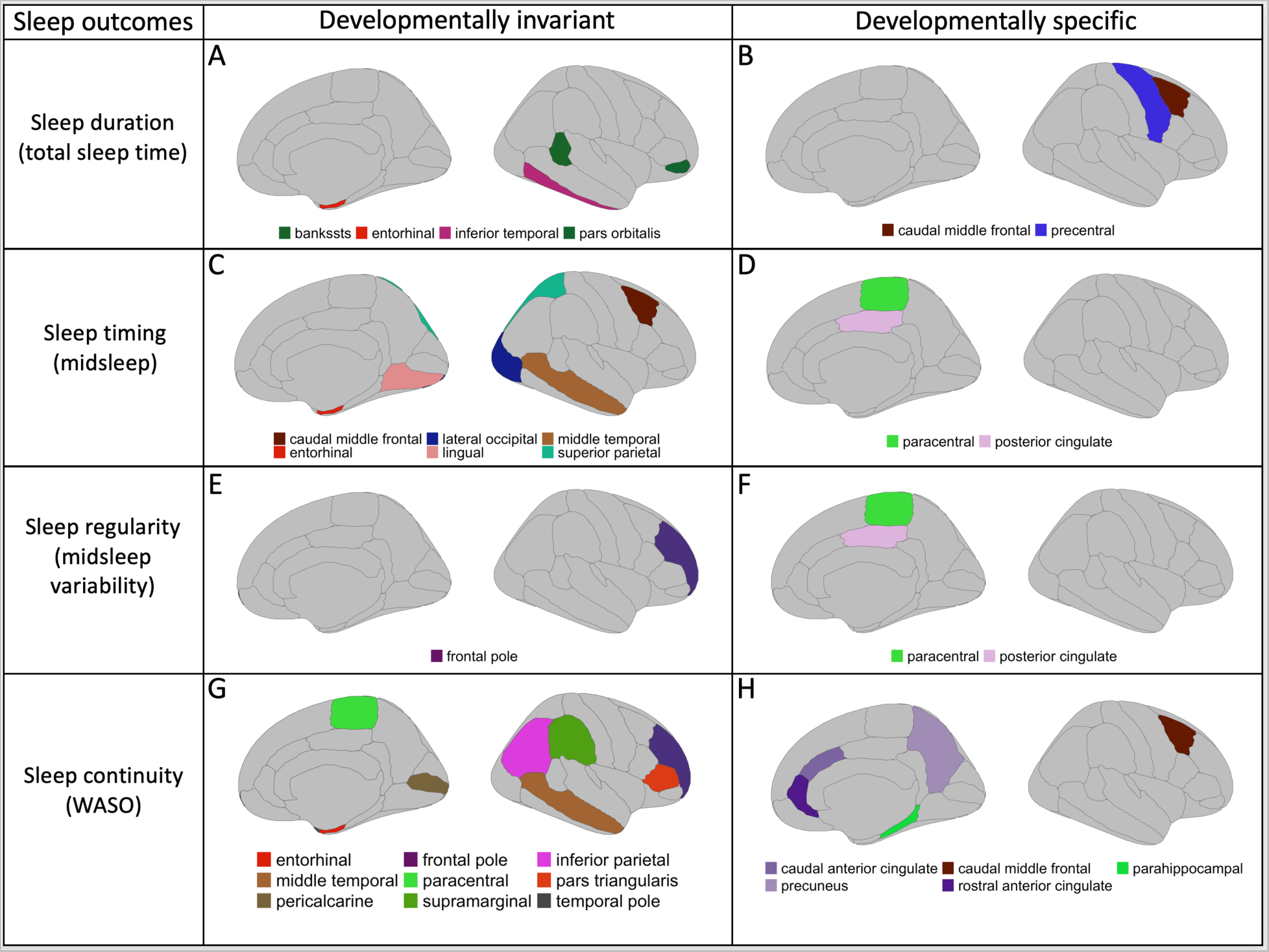
Developmentally invariant and specific relationships between sleep patterns and cortical gyrification. Figure 1 shows brain regions with developmentally invariant or specific relationships between sleep patterns and cortical gyrification. Panels A-B show sleep duration relationships, panels C-D show sleep timing relationships, panels E-F show sleep regularity relationships, and panels G-H show sleep continuity relationships. Each panel contains a legend with colors for the brain regions with non-zero coefficients identified with GLINTERNET.

**Table 2.**
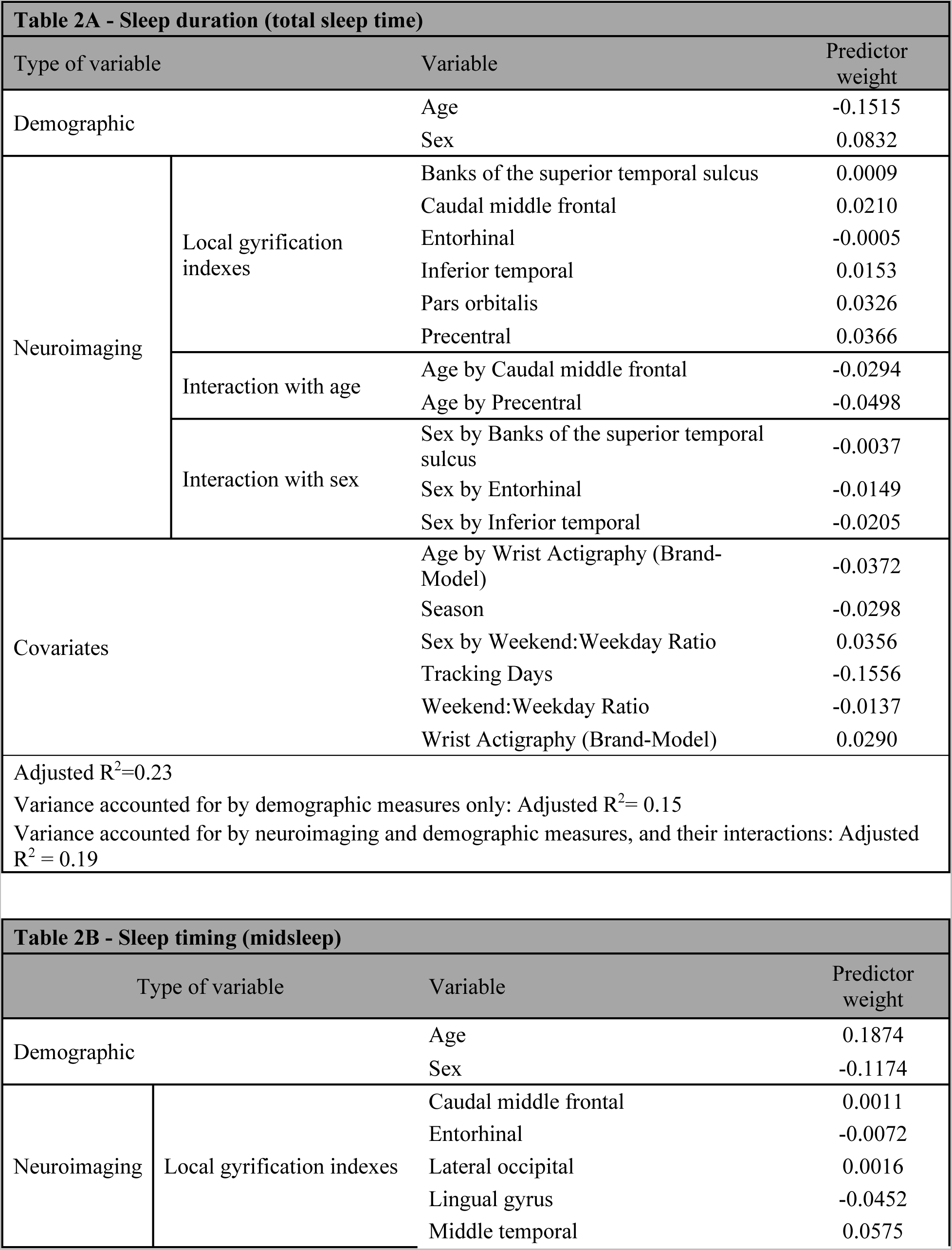
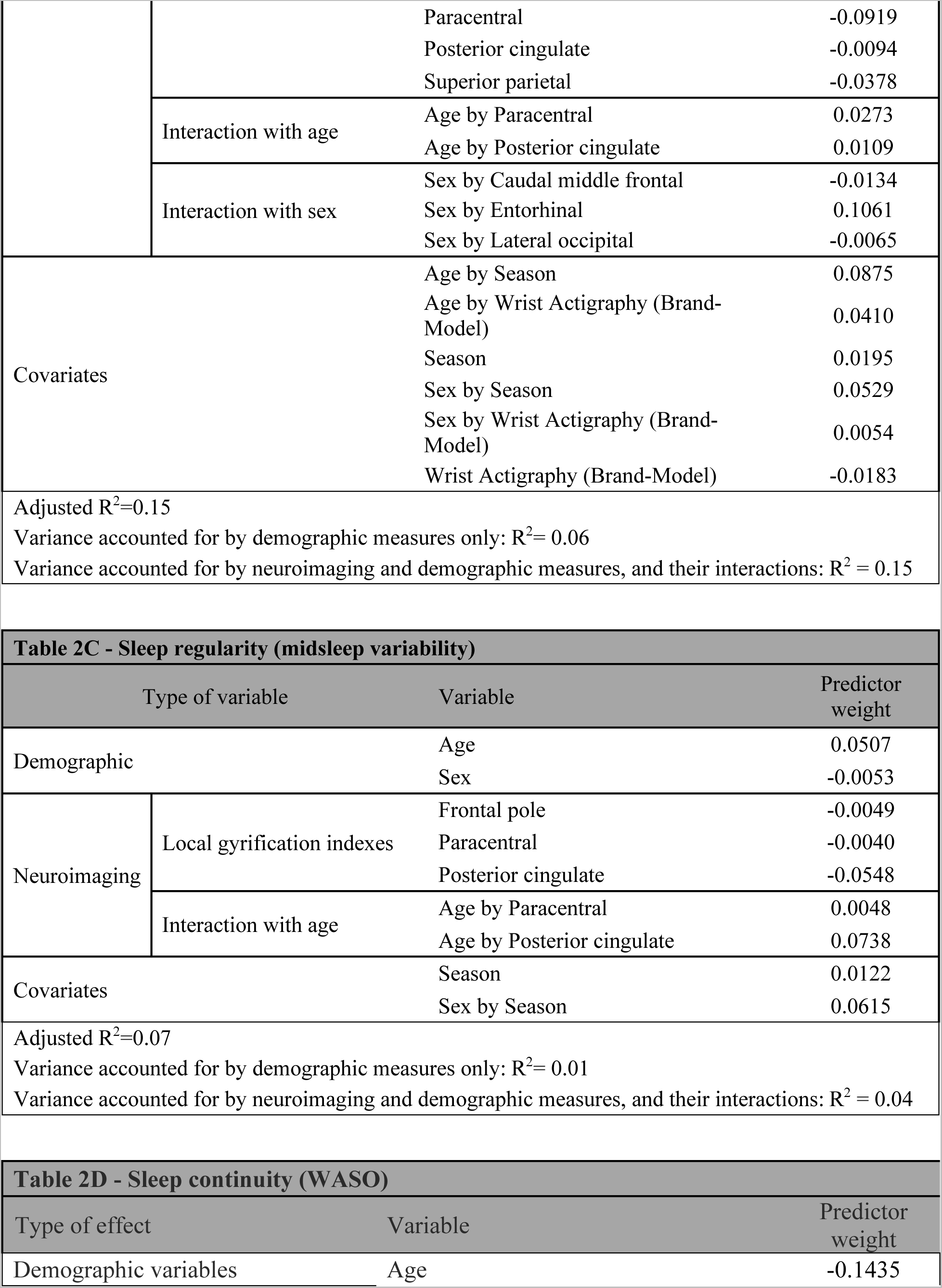
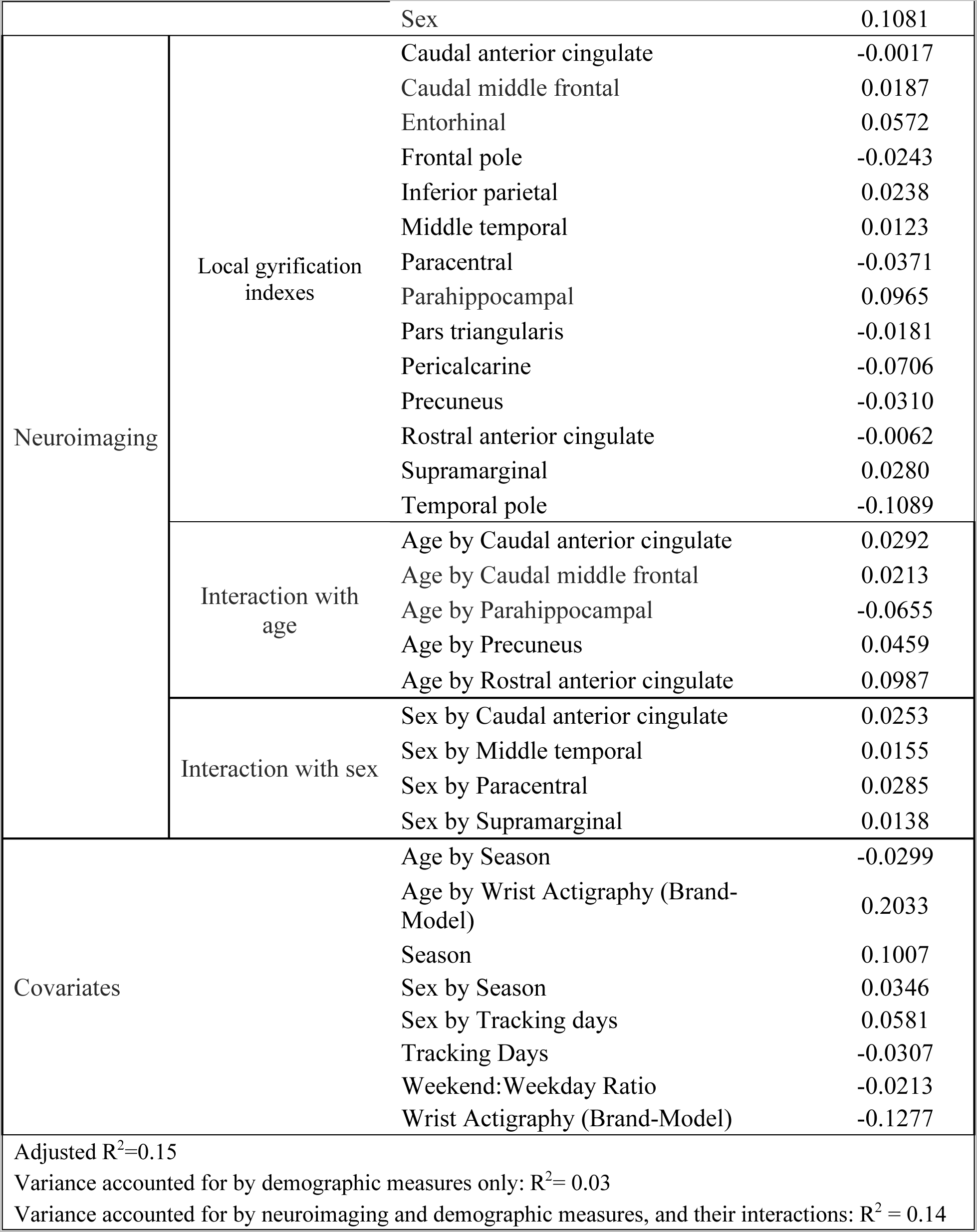
Main effects and interactions associated with sleep outcomes.

### Cortical Gyrification and Age

Overall, there was a significant negative linear association between age and cortical gyrification (P < 0.05).

### Sleep duration (total sleep time)

All features selected in this model explained 23% of the variability in sleep duration (Table 2A). Shorter sleep duration was associated with older age and males had shorter sleep duration in comparison to females.

We observed several developmentally invariant relationships between lGI and sleep duration in frontal and temporal regions. Greater lGI in frontal and temporal regions was associated with longer sleep duration. In addition, greater lGI in temporal regions was associated with longer sleep duration in male participants only. While the regularized regression model indicated that sex moderated the association between other temporal regions and sleep duration, this moderation effect was not significant in the traditional regression model (Supplemental Table 6).

We also found developmentally specific relationships between frontal lGI and sleep duration (Figure 2A), with Johnson-Neyman analyses showing that greater lGI was associated with longer sleep duration in participants 9-16 years old, but not in individuals 16-26 years old (Supplemental Table 7).

**Figure 2.**
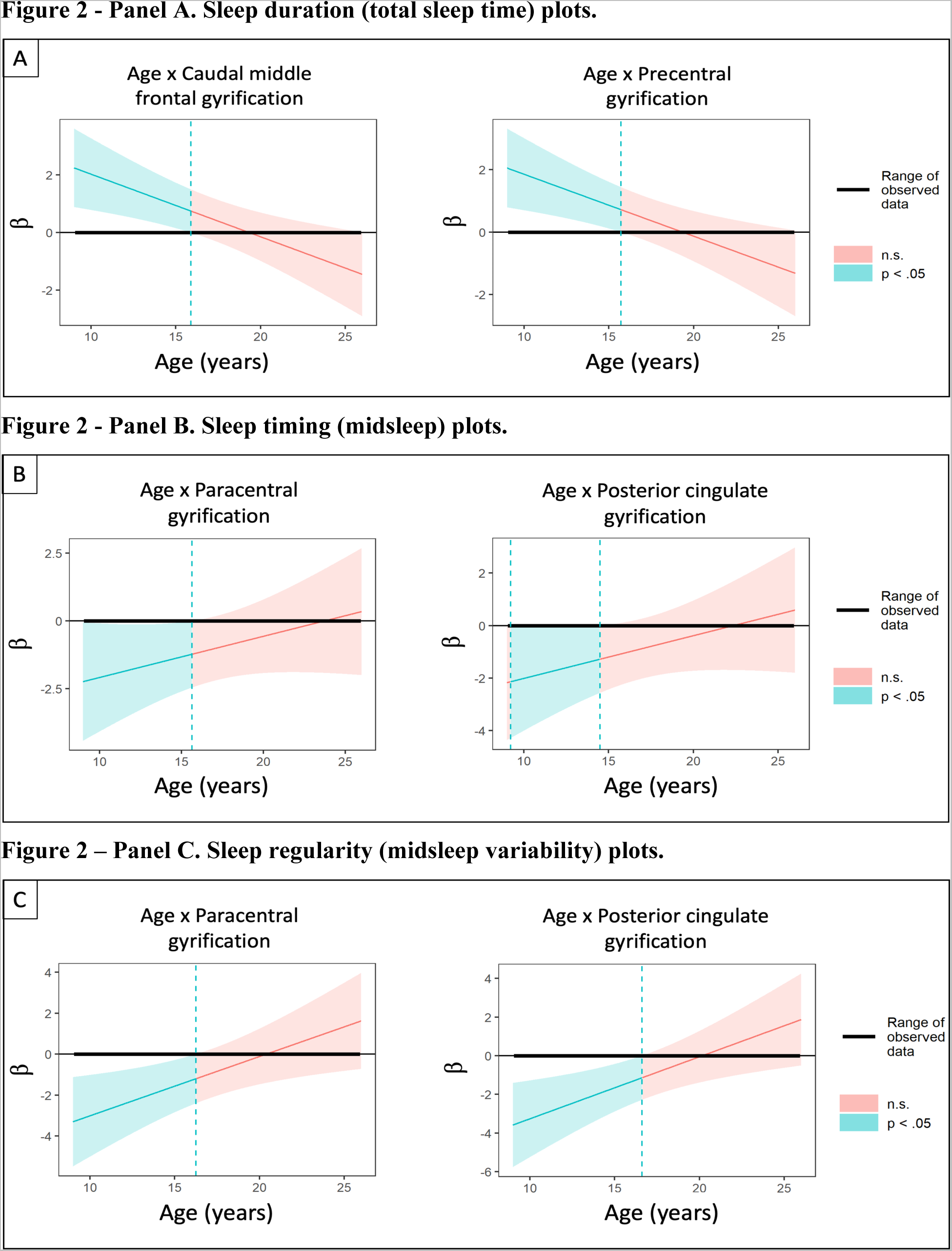
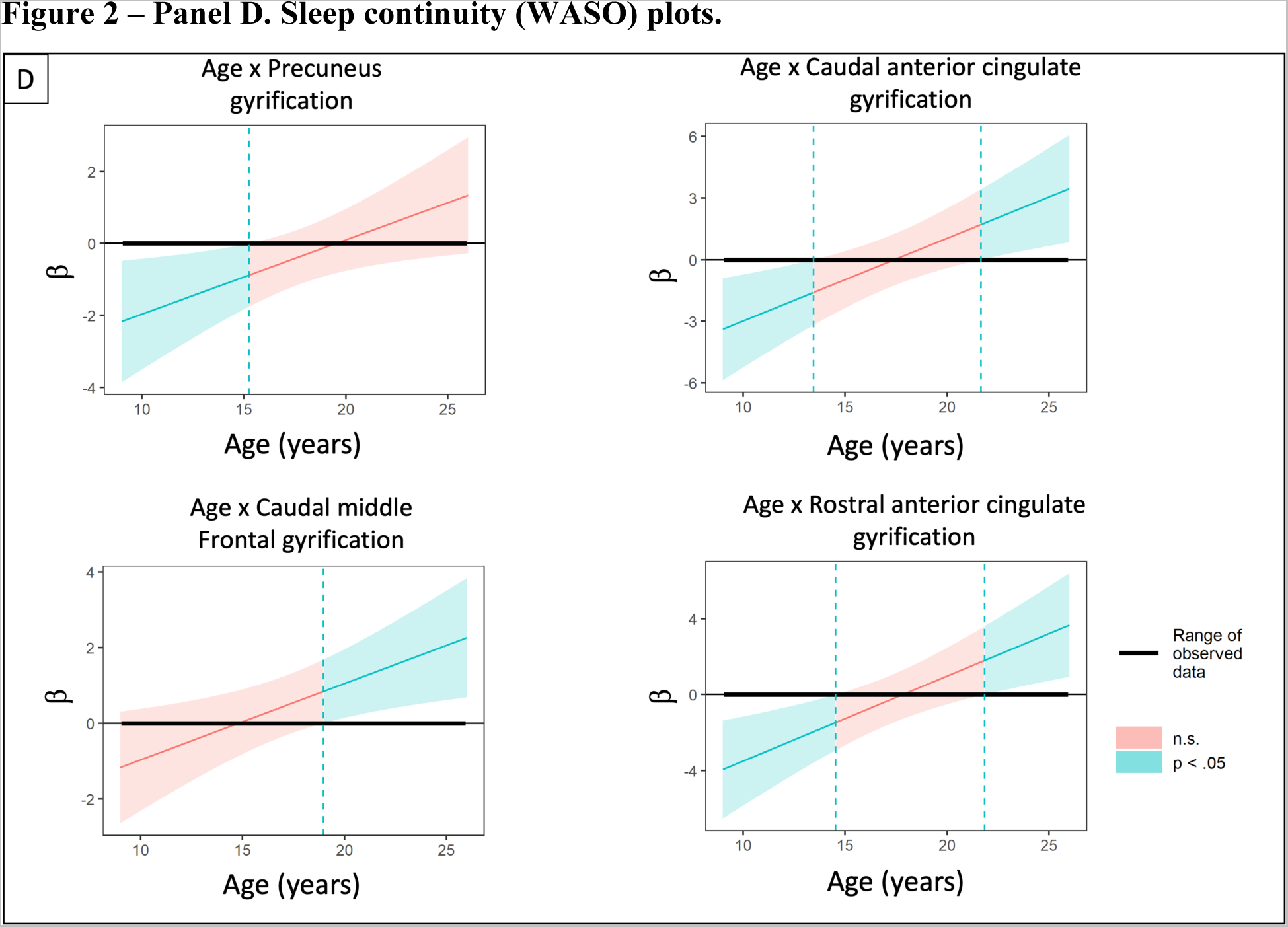
Johnson-Neyman plots for developmentally specific relationships. Figure 2 shows Johnson-Neyman plots for developmentally specific relationships (only during specific windows of development). Panel A shows sleep duration relationships, panel B shows sleep timing relationships, panel C shows sleep regularity relationships, and panel D shows sleep continuity relationships. In each plot, the X-axis shows the moderator (age - years) and the Y-axis shows the conditional association between sleep and cortical gyrification. Statistically significant relationships are shown in blue and non-significant relationships are shown in red.

### Sleep timing (midsleep)

Features selected in this model explained 15% of the variability in sleep timing (Table 2B). Later midsleep was associated with older age and being male.

We identified developmentally invariant relationships in frontal, temporal, parietal and occipital regions. Greater lGI in the lateral occipital and middle temporal regions was associated with later sleep timing, while greater lGI in the entorhinal, lingual and superior parietal regions was associated with earlier sleep timing. In female participants only, greater lGI in the frontal regions was associated with earlier sleep timing. In male participants only, greater lGI in temporal regions was associated with earlier sleep timing. While the regularized regression model indicated that sex moderated the association between occipital regions and sleep timing, this moderation effect was not significant in the regression model (Supplemental Table 6).

Developmentally specific relationships were observed in frontal and limbic regions (Figure 2B). Greater lGI in these regions was associated with earlier sleep timing in participants 9-15 years old, but not in older adolescents and young adults (16-26 years old; Supplemental Table 7).

### Sleep regularity (midsleep variability)

All features selected in this model explained 7% of the variability on sleep regularity (Table 2C). Greater variability in sleep regularity was associated with older age and being male.

A developmentally invariant relationship was identified in frontal regions. Greater lGI in this region was associated with lower variability in sleep regularity. There were no non-zero interactions between sex and lGI on sleep regularity.

Frontal and limbic regions showed developmentally specific relationships (Figure 2C). In these regions, greater lGI was associated with lower sleep variability in participants 9-16 years old, but not in individuals older than 16 (Supplemental Table 7).

### Sleep continuity (WASO)

Features selected in this model explained 15% of the variability in sleep continuity (Table 2D).

Several developmentally invariant relationships were identified. Greater lGI in the entorhinal, inferior parietal, middle temporal, and supramarginal regions was associated with longer WASO, while greater lGI in the frontal pole, paracentral, pars triangularis, pericalcarine, and temporal pole regions was associated with shorter WASO (i.e., better sleep continuity). In male participants only, greater lGI in the paracentral regions was associated with shorter WASO. While the group-lasso model indicated that sex moderated the association between WASO and caudal anterior cingulate, middle temporal, and supramarginal lGI, these moderation effects were not significant in the linear regression models (Supplemental Table 5).

Developmentally specific relationships were observed in five regions. In the precuneus, greater lGI was associated with shorter WASO in participants 9-16 years old only. In the caudal middle frontal regions, greater lGI was associated with greater WASO in participants 19-26 years old only. Interestingly, the anterior and caudal cingulate portions showed more complex relationships with sleep continuity: greater lGI was associated with shorter WASO in participants 9-14 years old and with longer WASO in participants 21-26 years old (Figure 2D; Supplemental Table 7).

## DISCUSSION

We examined to how measures of cortical gyrification were related to sleep characteristics and tested the extent to which these relationships were stable over age (developmentally invariant) or present only during specific windows of development (developmentally specific) in typically developing young people. Findings showed that cortical gyrification in diverse brain regions was associated with all investigated sleep characteristics. In most cases, greater cortical gyrification was associated with longer sleep duration, earlier sleep timing, lower variability in sleep regularity, and shorter time awake after sleep onset (characteristics generally associated with “better” sleep). Additionally, and in line with our hypotheses, developmentally specific associations were present and more common during late childhood through early-to-mid adolescence.

Across typically developing young people, increased cortical gyrification in a wide range of cortical regions was associated with characteristics generally associated with better sleep. These regions are involved in a number of functions including affective processing (frontal pole) [59], language processing (inferior temporal gyrus) [60, 61], memory (middle temporal gyrus) [62, 63], visual processing (inferior temporal gyrus) [60], and working memory (frontal pole) [59, 64]. In line with these findings, other macrostructural properties of the adolescent brain, such as volume and cortical thickness, have been associated with sleep [1, 65–67]. Some of these developmentally invariant relationships involved regions (e.g., middle frontal, supramarginal and parietal regions) commonly associated with the frontoparietal network (FPN), a brain network implicated with executive control and complex cognitive functions [68]. Previous studies have shown that poor sleep during brain development is detrimental to FPN function [69] and connectivity between FPN and other networks (e.g., limbic network) [70]. However, no study has shown a link between cortical gyrification of regions within the FPN and sleep characteristics, and our findings show that sleep is important for this network throughout all stages of brain development. Considering that cortical gyrification has been associated with age- and cognition-related decline later in life [10, 15], our findings suggest that better sleep may be an important factor for the brain-cognition relationships across lifespan and that maintaining better sleep since early age may promote better cognitive functioning later in life.

Our findings also add to an existing literature demonstrating that cortical gyrification across lifespan may be associated with modifiable lifestyle behaviors, such as physical activity [71], dietary patterns [72], and substance use (e.g., tobacco, and alcohol) [71]. For example, in one of these studies [71], a composite “lifestyle risk” factor was calculated based upon alcohol consumption, smoking, physical activity, and social integration in older adults. They demonstrated that more optimal health behaviors were associated with increased cortical gyrification (which is typically viewed as “healthier”) and decelerated decline in multiple brain regions [71]. Given that, overall, gyrification has shown a positive relationship with cognitive functions in adults [10, 15], these findings suggest a potential mechanism by which lifestyle factors may impact these functions later in life. The multiple dimensions of sleep studied here - such as sleep duration, timing, regularity, and continuity - are amenable to efficacious behavioral interventions, providing an avenue for understanding and promoting healthy development. The period of adolescence and young adulthood is particularly important because of its foundational role in adaptive functioning during adulthood. Our findings of the association between gyrification and sleep characteristics provide clues to optimizing brain development. Future studies evaluating the effect of lifestyle risk factors on cortical gyrification, or other macrostructural properties of the brain should also consider the role of sleep.

In addition, for some of these developmentally invariant relationships, sex played a moderating role, with males showing stronger relationships between cortical gyrification and sleep than females in areas implicated with memory, visual and language processing. Sex differences play an important role in many neural and behavioral functions in typically developing young people, and these differences have been linked to the cascade of maturational changes (e.g., physical, psychological, and social) happening during this period [73, 74]. Previous studies have shown sex differences in sleep physiology [75], secondary clinical outcomes of sleep problems [76], and cortical gyrification of typically developing young people [77]. Our findings build upon the existing literature to show that sex differences are also present in the relationship between sleep and brain and suggest that cortical gyrification could be a structural component that contributing to sex differences found in the relationship between sleep and cognition [78, 79].

Increased cortical gyrification was associated with more optimal sleep in late childhood through early-to-mid adolescence. In line with these findings, we have recently shown that other macrostructural properties of the brain (e.g., subcortical volume and cortical thickness) also present developmentally specific relationships with sleep from late childhood to mid-adolescence [1], reinforcing the importance of sleep during this period of brain development. The transition from childhood to adulthood represents a period of development marked by several physical, cognitive, and social changes, with early-to-mid adolescence being particularly associated with important maturational changes in the organization of brain networks [80, 81]. Studies have shown that abnormalities in this tuning process are associated with mental health issues later in life [82]. Our findings showed a developmentally specific relationship in frontal-parietal-limbic brain regions, including those typically associated with the default mode network (DMN; e.g., anterior cingulate, posterior cingulate, precuneus). The DMN is implicated with emotional and episodic memory processing and undergoes important strengthening of functional connectivity between its regions during late childhood through early-to-mid adolescence (7-15 years old) [83–85]. In addition, previous studies have shown that more optimal sleep is associated with increased network connectivity in DMN regions of adolescents and young adults [86, 87]. Our findings show that sleep patterns are strongly associated with regions associated with DMN, particularly during an important period for their development. More importantly, they also suggest early adolescence as a window of vulnerability for the effects of less optimal sleep and as a window of opportunity for early interventions aiming to improve trajectories for positive development.

There were limitations in this study. While this study raised important questions relevant to the relationship between sleep and brain, future prospective longitudinal studies would benefit from even bigger samples that included larger age ranges, different populations at risk, more racial and ethnic diversity and other possible individual factors affecting this relationship during brain development (e.g., pubertal development, sex hormones, socioeconomic status). Although group-lasso is a strong feature selection method that can identify even small effects important for the for tested models, non-zero coefficients identified with these methods do not always translate to significance in standard regression models. While LASSO regularization includes all potential variables and uses a shrinkage parameter to reduce the coefficient of non-relevant variables toward zero, standard regression models test a null hypothesis using pre-selected variables. Therefore, future studies using methods other than standard regression are needed to further explore non-zero coefficients. In addition, LASSO reduces the coefficients of highly correlated variables, leading to the possible exclusion of gyrification measures that are associated with sleep behavior (and correlated with other selected variability). Given that cortical gyrification measures show high correlation (Supplemental Figure 1), in the future, we may want to consider additional methods to best account for this covariance. Finally, this study explored linear relationships between sleep and cortical gyrification. Future studies should also explore potential non-linear relationships since many changes during adolescence follow both linear and non-linear trajectories.

Cortical gyrification remains an understudied neural correlate of sleep patterns, despite its ability to provide unique information about brain morphology, particularly during brain development. Given that cortical gyrification naturally declines over the lifespan, our findings may indicate that sleep patterns during childhood and early adolescence have the potential to accelerate or decelerate multiple neurodevelopmental processes (e.g., cognitive and emotional). Future work should evaluate these relationships within person, particularly during early-to-mid adolescence, to better understand the association between individual patterns and cortical gyrification. Furthermore, considering that aberrant cortical gyrification has been associated with cognitive decline and psychiatric disorders later in life, our findings further suggest that better sleep may be a protective factor for age-related effects on the brain since early age and especially in early-mid adolescence, a developmental period of high vulnerability for symptom escalation and high opportunity for intervention.

## Supplemental Materials

### METHODS

#### Participants

Participant consent or assent was collected at enrollment for each individual study included in NAPS and permitted sharing of de-identified data. Studies were considered for inclusion in NAPS if they included: (1) baseline actigraphic sleep monitoring reflecting naturalistic sleep; (2) a sMRI scan; and (3) participants aged 8.0–30.9 years-old (inclusive). Participant-level inclusion criteria were: (1) 9.0–25.9 years-old; (2) absence of current psychiatric diagnosis based on clinical interview (i.e. KSADS, SCID); (3) no current psychotropic or hypnotic medication use; (4) ≥5 days of good quality actigraphic sleep monitoring composed of both weekday and weekend days; (5) good quality MRI scan.

#### Sleep data

Trained scorers blinded to neuroimaging data manually identified rest intervals based on a combination of event markers indicated by participants and clear changes in activity and (if available) environmental light level recorded by the device. Brand-specific sleep scoring algorithms estimated sleep within each rest interval [1–6]. We implemented additional semi-automated quality assurance procedures using in-house R scripts, including identification of the main rest interval (defined as the longest rest interval each day), removal of invalid sleep intervals containing ≥1 hour of off-wrist time or recording errors [2, 7], time adjustment for daylight savings time, and final visual inspection of sleep intervals on raster plots.

### SUPPLEMENTAL TABLES

**Supplemental Table 1.**
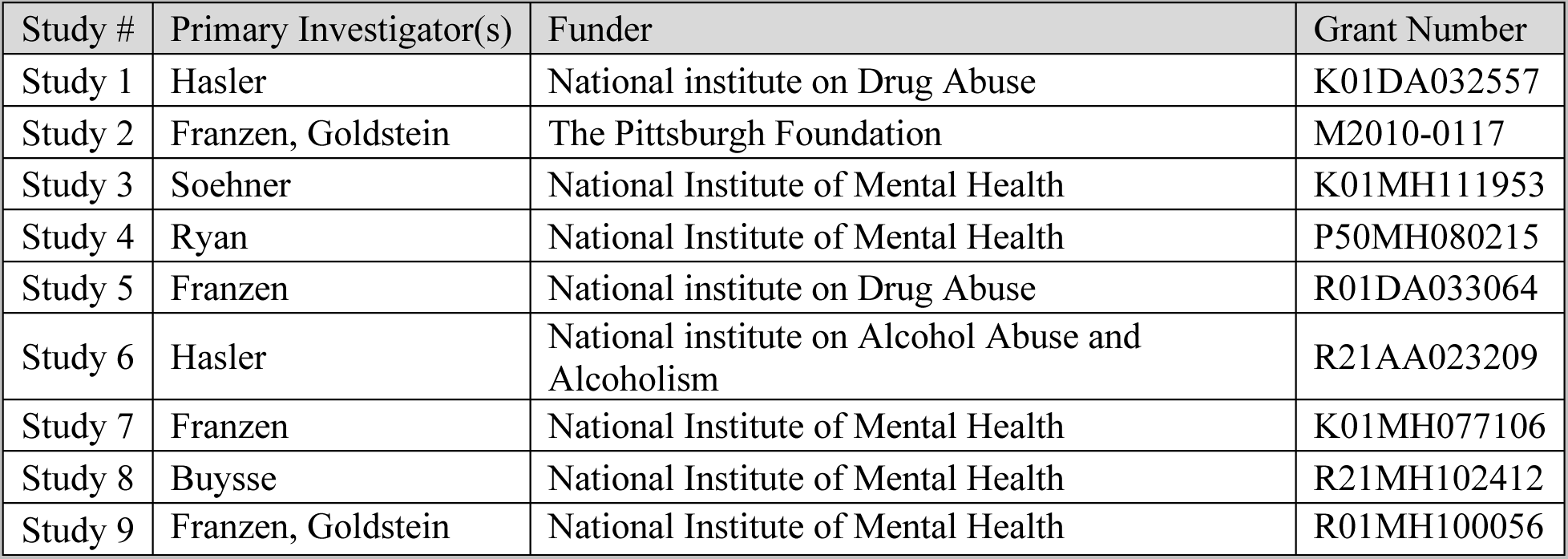
Studies included in the Neuroimaging and Pediatric Sleep (NAPS) databank.

**Supplemental Table 2.** Actigraphy characteristics by study.

| Study | Enrollment Year (Min-Max) | Actigraph (Brand-Model) |
| --- | --- | --- |
| Study 1 | 2013-2017 | Philips-Spectrum |
| Study 2 | 2012-2013 | Philips-Actiwatch2 |
| Study 3 | 2017-2019 | Philips-Spectrum |
| Study 4 | 2009-2013 | AMI-Motionlogger |
| Study 5 | 2012-2014 | Philips-Actiwatch2 |
| Study 6 | 2014-2017 | Philips-Spectrum |
| Study 7 | 2009-2012 | Philips/MiniMitter-Actiwatch64 |
| Study 8 | 2014-2016 | Philips-Actiwatch2 |
| Study 9 | 2017-2019 | Phillips-Spectrum Series |

**Supplemental Table 3.** Acquisition protocols.

| Study | Scanner Manufacturer | Tesla Strength | Scanner Model | Repetition Time (ms) | Echo Time (ms) | Inversion Time (ms) | Flip Angle | Voxel Size (mm) | Phase Field of View |
| --- | --- | --- | --- | --- | --- | --- | --- | --- | --- |
| Study 1 | Siemens | 3 | Trio Tim | 2.2 | 0.00331 | 1 | 9 | 1 | 75 |
| Study 2 | Siemens | 3 | Biograph mMR | 2.2 | 0.00335 | 1 | 9 | 1 | 75 |
| Study 3 | Siemens | 3 | Trio Tim | 2.2 | 0.00331 | 1 | 9 | 1 | 75 |
| Study 4 | Siemens | 3 | Prisma | 5 | 0.00297 | .7/2.5 | 4/5 | 1 | 93.8 |
| Study 5 | Siemens | 3 | Trio Tim | 2.1 | 0.00343 | 1.05 | 8 | 1 | 81.25 |
| Study 6 | Siemens | 3 | Trio Tim | 2.2 | 0.00331 | 1 | 9 | 1 | 75 |
| Study 7 | Siemens | 3 | Trio Tim | 1.5 | 0.00319 | 0.8 | 8 | 1 | 100 |
| Study 8 | Siemens | 3 | Trio Tim | 2.2 | 0.00331 | 1 | 9 | 1 | 75 |
| Study 9 | Siemens | 3 | Trio Tim | 2.1 | 0.00331 | 1.05 | 8 | 1 | 81.25 |
| Study 1 | Siemens | 3 | Prisma | 5 | 0.00297 | 1.05 | 8 | 1 | 93.75 |

Abbreviations: ms = milliseconds; mm = millimeter

**Supplemental Table 4.**
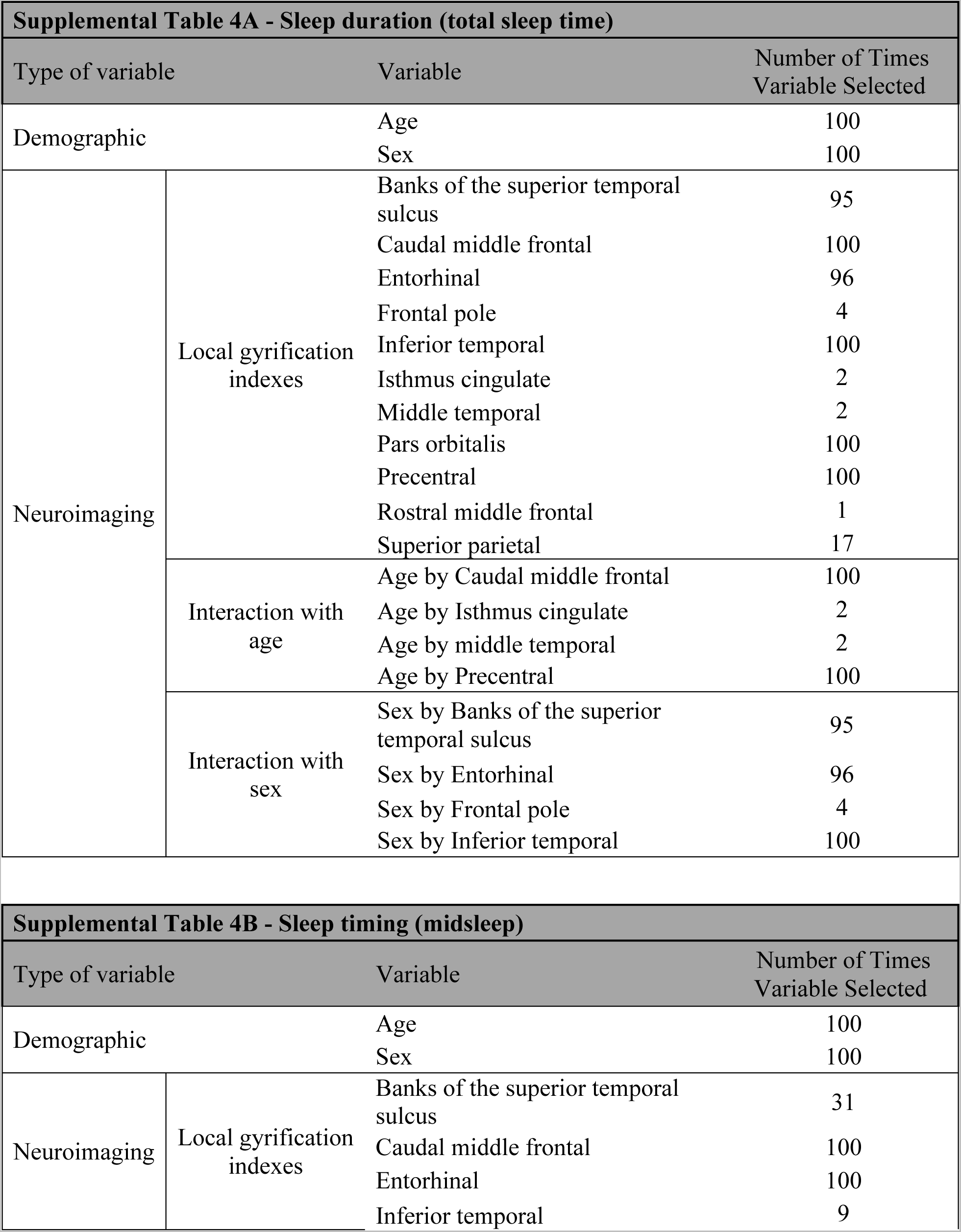
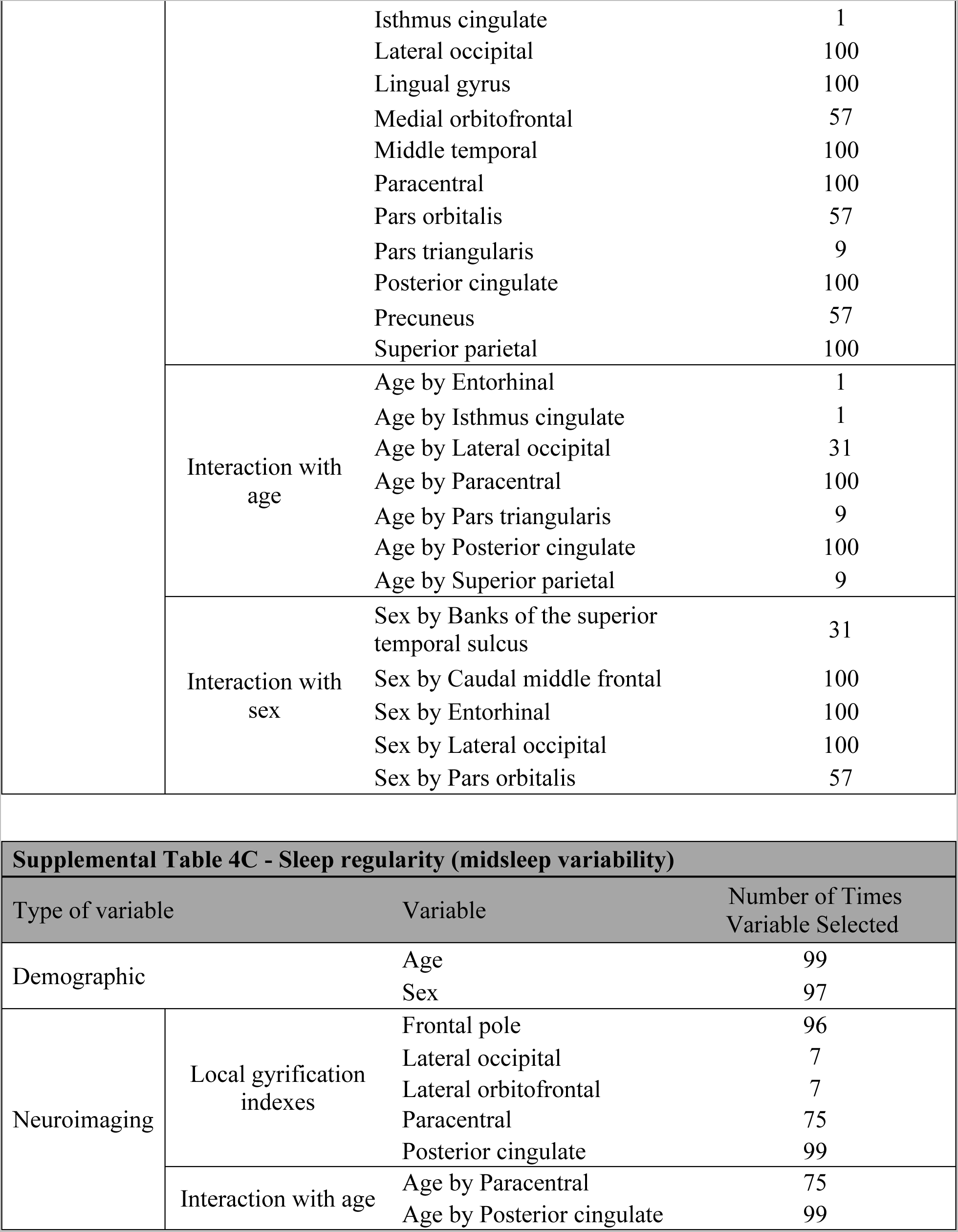
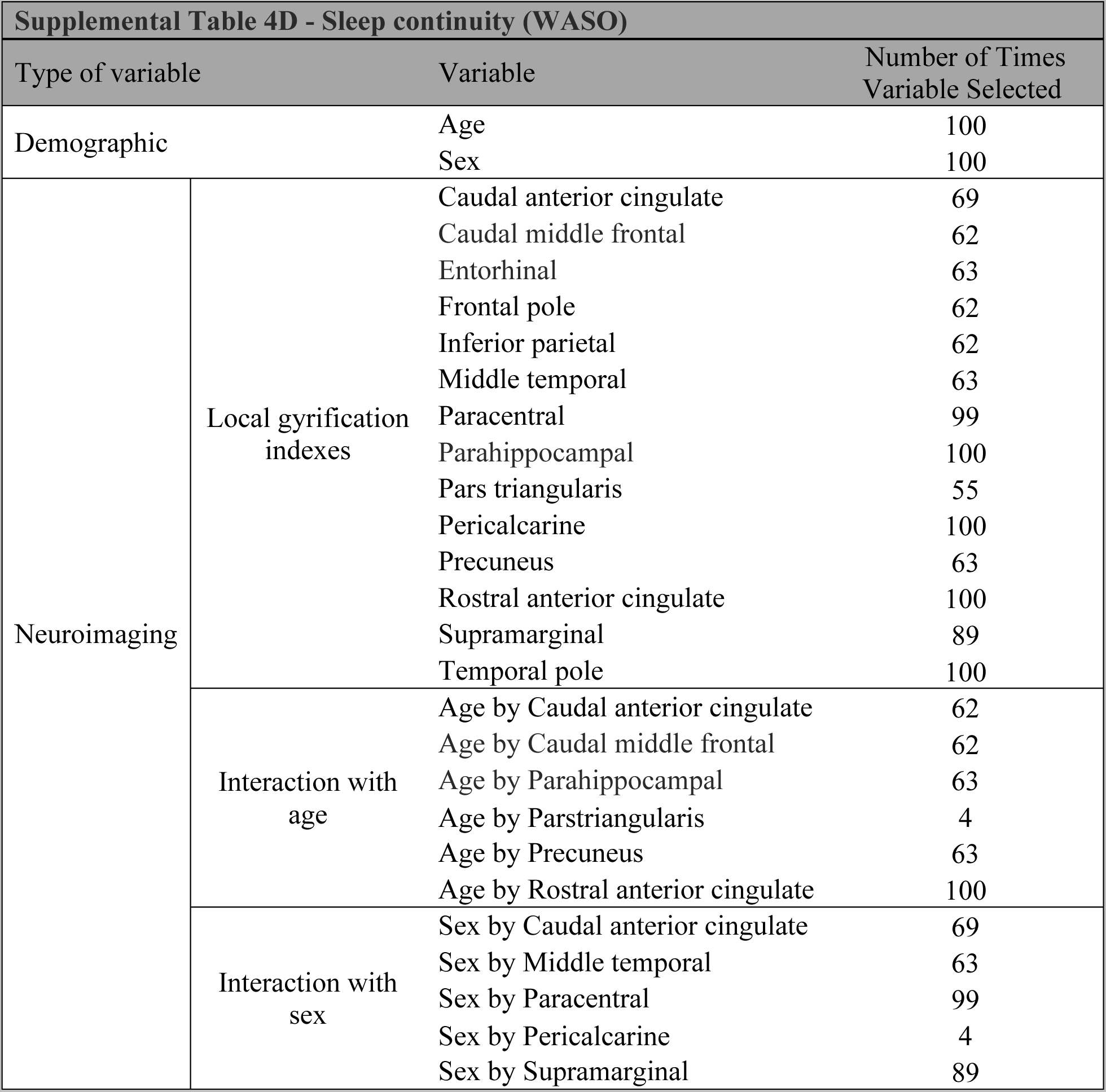
Stability of non-zero demographic and neuroimaging predictors that were selected in 100 group-lasso iterations.

**Supplemental Table 5.** Multiple regression models.

| Models | F | Adjusted R-squared | P-value |
| --- | --- | --- | --- |
| Sleep duration (total sleep time) | 4.94 | 0.23 | <0.001 |
| Sleep timing (midsleep) | 3.08 | 0.15 | <0.001 |
| Sleep regularity (midsleep variability) | 2.92 | 0.07 | 0.002 |
| Sleep continuity (WASO) | 2.31 | 0.15 | <0.001 |

**Supplemental Table 6.**
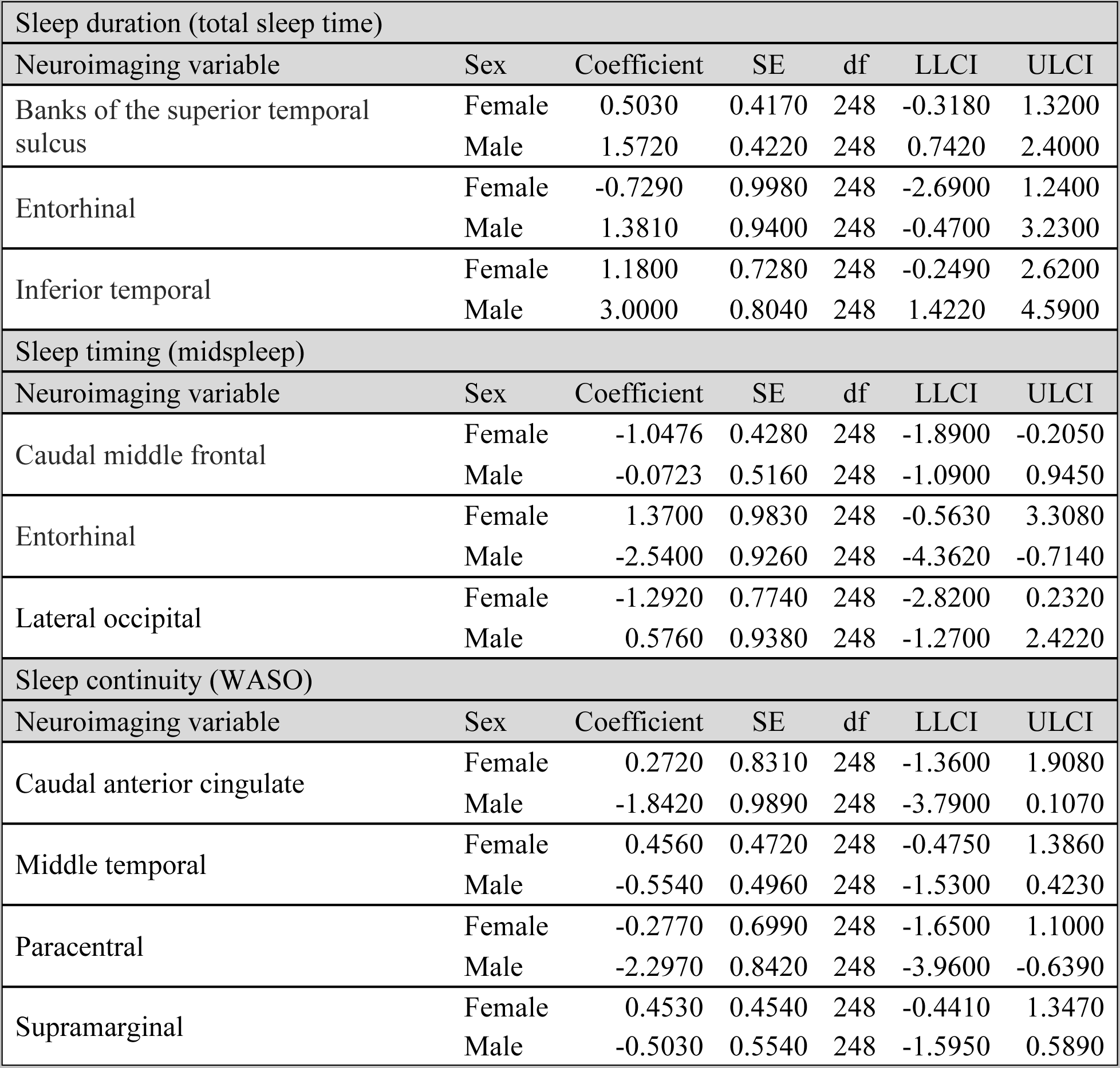
Interaction effects between sex and gyrification indexes.

**Supplemental Table 7.** Interaction effects between age and gyrification indexes.

| Sleep duration (total sleep time) |  |
| --- | --- |
| Neuroimaging variable | JN interval (years) |
| Caudal middle frontal | < 15.9 |
| Precentral | < 15.7 |
| Sleep timing (midsleep) |  |
| Neuroimaging variable | JN interval (years) |
| Paracentral | < 15.7 |
| Posterior cingulate | < 14.5 |
| Sleep regularity (midsleep variability) |  |
| Neuroimaging variable | JN interval (years) |
| Paracentral | < 16.3 |
| Posterior cingulate | < 16.6 |
| Sleep continuity (WASO) |  |
| Neuroimaging variable | JN interval (years) |
| Caudal anterior cingulate | < 13.5 and > 21.6 |
| Caudal middle frontal | > 18.9 |
| Parahippocampal | - |
| Precuneus | < 15.2 |
| Rostral anterior cingulate | < 14.5 and > 21.9 |

Abbreviations: JN = Johnson-Neyman.

### SUPPLEMENTAL FIGURES

**Supplemental Figure 1.**
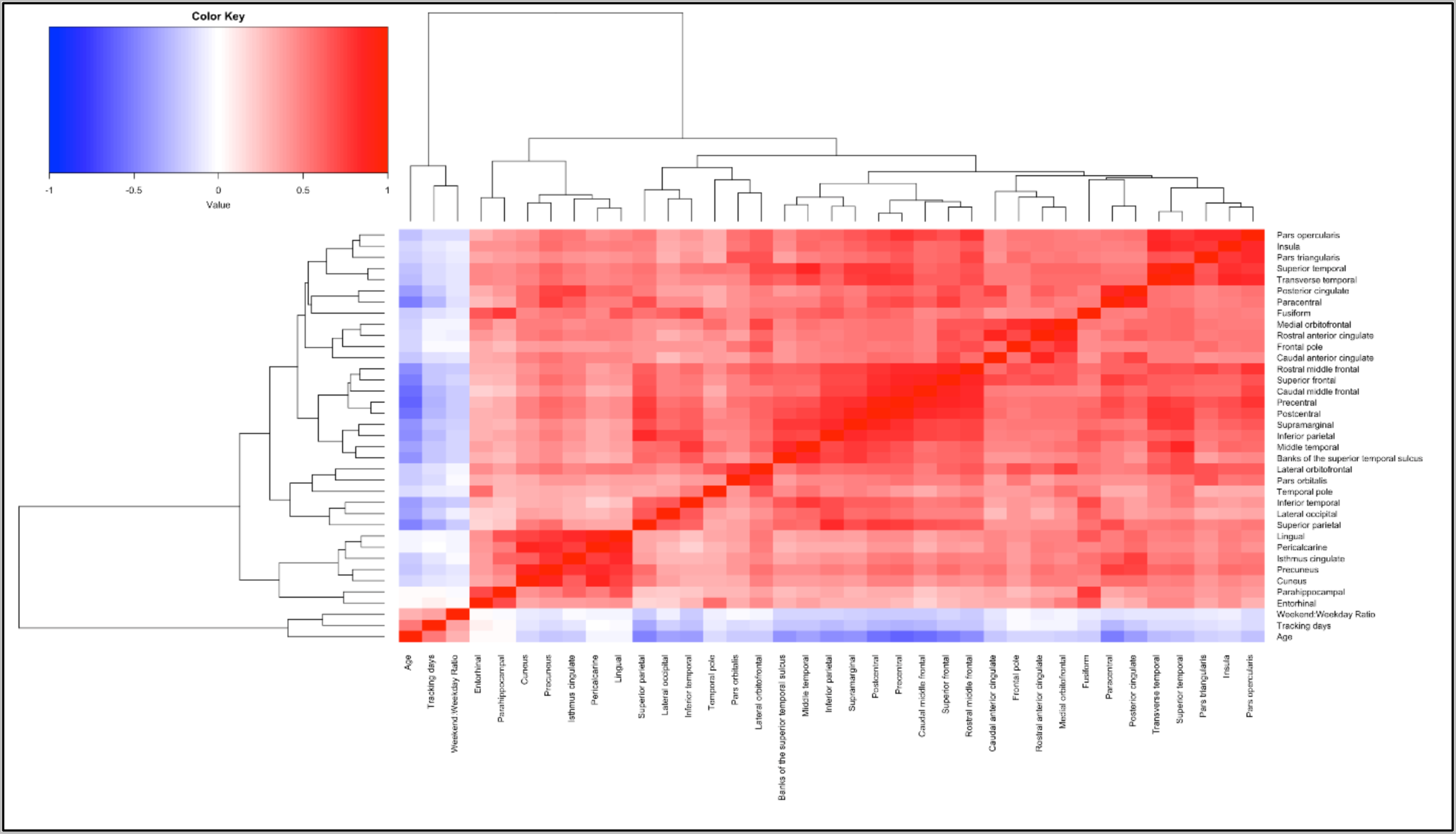
Correlation heatmap for cortical gyrification measures. Supplemental figure 1 depicts a heatmap showing pair correlations across continuous variables included in the brain-sleep models. Blue color indicates negative correlations and red color indicates positive correlations.

